# Yeast replicative aging leads to permanent cell cycle arrest in G1 effectuated by the start repressor Whi5

**DOI:** 10.1101/353664

**Authors:** Jing Yang, Ziwei Wang, Xili Liu, Hao Li, Qi Ouyang

## Abstract

Yeast replicative aging has been a canonical model for aging research. Since replicative aging eventually leads to permanent cell cycle arrest, a fundamental question is how cells sense the signals from aging and communicate that to the cell cycle control machineries. Using microfluidic devices to track individual mother cells labeled by two different cell cycle markers Whi5-tdTomato and Myo1-EGFP, we measured the length of different cell cycle phases as a function of age and the distribution of cell death in different cell cycle phases. We found that the majority of the cells died in the G1 phase, and their G1 cell cycle length increased drastically in the last few cell divisions. This increase of G1 length correlates with the increase of the nuclear concentration of Whi5, which is a major transcriptional suppressor of the cell cycle start check point. Interestingly, this correlation is apparent only above a threshold concentration of Whi5. We show that in response to external stress, Whi5 concentration increases and cell growth slows down in a Whi5 dependent manner, and that Whi5 deletion significantly extends the lifespan. Together these data suggest the existence of a programmed control to arrest cell cycle in G1 in response to stress signals due to aging, and that Whi5 is a major mediator of this process. Our findings may have important implications in understanding senescence and cancer in mammalian cells, which have a parallel G1/S control system with Rb (a well known tumor suppressor) as the analog of Whi5.

**Significance statement:** In this work, we used microfluidic devices to track individual mother cells labeled by two cell cycle markers Whi5-tdTomato and Myo1-EGFP. We found that aging leads to significant lengthening of G1 phase in old cells and the eventual permanent cell cycle arrest in G1, and Whi5 plays an important role in implementing such a program. We show that oxidative stress can lead to the increase of Whi5 expression and the slow-down of cell division. Furthermore, Whi5 deletion significantly extends the lifespan. The result suggest the existence of a programmed control to arrest cell cycle in G1 in response to stress signals due to aging, and that Whi5 is a major mediator of this process.

## Introduction

The replicative aging of the budding yeast *Saccharomyces cerevisiae* has served as a canonical model for aging research and has led to the discovery of conserved longevity pathways across species (Johnson, Sinclair et al. 1999, Kenyon 2001, Rogina and Helfand 2004, Kaeberlein 2010). The replicative lifespan of a mother cell is defined as the number of daughter cells produced by the mother before permanent cell cycle arrest; thus replicative aging is ultimately tied to cell cycle regulation. Previous studies have shown that as the cells grow old, their cell size progressively increases and their cell cycle significantly lengthens for the last few cell divisions (Egilmez and Jazwinski 1989, Xie, Zhang et al. 2012), and that cells die with different terminal morphologies indicative of different cell cycle phases (Xie, Zhang et al. 2012, Delaney, Chou et al. 2013). However, it remains largely unknown how the status of aging is sensed and communicated to the cell cycle control machineries to eventually stop the cell cycle permanently.

Budding yeast cell cycle is composed of four distinct phases: the first gap phase (G1), DNA synthesis phase (S), the second gap phase (G2), and mitosis (M). Cells make irreversible cell cycle commitment at the end of G1 phase after passing the Start checkpoint, accompanied by the transcriptional activation of ~200 G1-specific genes (Eser, Falleur-Fettig et al. 2011). Prior to the Start, cells are sensitive to G1 arrest. The periodic transcription at Start depends on two transcription factor complexes: SBF (Swi4p-Swi6p complex) and MBF (Mbp1-Swi6 complex) (Breeden 2003), and they are transcriptionally repressed by the Start repressor, Whi5 (Costanzo, Nishikawa et al. 2004). Events at Start depend on the activation of Cdc28, which is activated by the upstream G1 cyclin Cln3 and then by the downstream G1 cyclins Cln1 and Cln2 (Cross 1995). Cdc28-Cln3 inside nucleus phosphorylates Whi5, moving it out of nucleus and thus partially release SBF and MBF, and as a result, Cln1 and Cln2 are transcribed. Cdc28-Cln1 and Cdc28-Cln2 further phosphorylate Whi5, establishing a positive feedback loop that leads to the sharp G1-S transition (Skotheim, Di Talia et al. 2008). Thus the transcriptional repressor Whi5, whose phosphorylation and re-localization is rate-limiting, is a key regulator of Start. Whi5 is localized in nucleus from the end of mitosis until the Start (Costanzo, Nishikawa et al. 2004, de Bruin, McDonald et al. 2004), and its nuclear localization has been used to mark the G1 phase (Ball, Marchand et al. 2011).

Given the observations that aged cells have prolonged cell cycle in their last few divisions and many permanently arrest their cell cycle with a terminal morphology suggestive of G1 (Johnston 1966, Egilmez and Jazwinski 1989, Xie, Zhang et al. 2012), it will be very informative to analyze the cell cycle dynamics as a function of age quantitatively with a focus on cell cycle regulators involved in controlling G1/S transition. This has not been feasible with the traditional micro-dissection technique, which is laborious and time consuming, and does not have sufficient optical resolution to follow molecular markers inside the mother cells.

An important aspect of cellular response closely tied to cell cycle control and aging is the stress response. It is known that in cell cultures dominated by young cells, the transcriptional regulators Msn2/4 are activated and turn on hundreds of genes in respond to general environmental stresses, and cells typically arrest in G1 phase (Shackelford, Kaufmann et al. 2000, Causton, Ren et al. 2001, Chang, Tseng et al. 2017). In the context of single cell aging, we have observed that Msn2/4 activity (based on a transcriptional reporter) progressively increases starting from the middle age (Xie, Zhang et al. 2012). These observations suggest some similarities between stress response and aging, but how they are linked mechanistically is not understood.

In this work, we analyze the interplay between aging, cell cycle control and stress response. We used a microfluidic system to track mother cells and follow cell cycle markers throughout their lifespan (Xie, Zhang et al. 2012, Zhang, Luo et al. 2012). We labeled mother cells by two different cell cycle fluorescent markers: Whi5-tdTomato and Myo1-EGFP. Whi5 is a transcriptional repressor that moves into the nucleus at the end of mitosis and moves out of the nucleus at the end of G1, and thus is used to mark G1 phase. Myo1 is an actomyosin ring component that forms a ring at the bud neck and the ring disappears after cytokinesis is completed; we used it to mark the end of M phase. Using this labeled strain and the microfluidic device, we simultaneously measured the lifespan, cell cycle dynamics, and Whi5 expression/localization in single mother cells throughout their lifespan. We found that the majority of the cells died in G1 phase, and their lifespan has a bell shaped distribution with a characteristic age. In contrast, cells died in M phase has a flat distribution. This suggests that G1 arrest is the general end point for cells died due to normal aging. We observed a progressive increase of Whi5 nuclear concentration from middle age and significant increase of cell cycle length in the last few cell divisions. For cells died in G1, the lengthening of their cell cycle is mainly attributed to the lengthening of G1, which correlates with Whi5 nuclear concentration. We found the exogenous oxidative stress increases Whi5 concentration and slows down cell growth in a Whi5 dependent manner, and that deletion of *WHI5* leads to significant lifespan extension. Combining these observations, we postulate that aging generates stresses and in response cell increases Whi5 nuclear concentration. After Whi5 concentration reached a threshold, it starts to slow down G1 and eventually stops cell cycle permanently in G1. Our study may have implications in understanding senescence in mammalian cells with a parallel G1/S control system, in which Rb (a well-known tumor repressor) is the analog of Whi5.

## Results

### Cell death in different cell cycle phases and the corresponding lifespan distributions

We constructed a yeast strain with two cell cycle fluorescent markers, tdTomato tagged Whi5 and GFP tagged Myo1, to characterize cell division in budding yeast (Fig. 1B). Whi5 is a transcriptional repressor that controls G1/S transition. It is imported to nucleus at the end of mitosis and stays in nucleus until the START, and the duration of Whi5’s stay in the nucleus has been used to measure the length of G1 phase (Ball, Marchand et al. 2011, Liu, Wang et al. 2015). Myo1 is a subunit of the type II myosin heavy chain that plays a critical role in cytokinesis and cell separation. It forms a ring at the incipient bud site in late G1 or early S phases, before bud emergence. This ring remains at the bud neck and disappears after cytokinesis is completed (Bi, Maddox et al. 1998). This marker is used to define the end of the M phase. Using this strain and the microfluidic devices we developed (Xie, Zhang et al. 2012, Zhang, Luo et al. 2012, Zou, Ren et al. 2017), we tracked single yeast mother cells throughout their lifespan by time-lapsed microscopy and quantified cell cycle dynamics and marker expression as a function of their replicative age.

We first quantified cell death in different cell cycle phases. Previously it was observed that dead cells assume different terminal morphologies suggestive of cell cycle arrest at different cell cycle phases (Xie, Zhang et al. 2012, Delaney, Chou et al. 2013). E.g., an un-budded terminal morphology is suggestive of G1 arrest while a large-budded morphology is indicative of arrest in the M phase. However, without cell cycle molecular markers, it is difficult to accurately determine the cell cycle phase at which a cell died. With the two molecular markers, we classified cell death as: 1) G1-death if whi5 is in the nucleus; 2) M-death if Whi5 is out of nucleus, the mother cell is elongated with a large bud, and the myo1 ring is still present (before the completion of cytokinesis); 3) S-death for the rest of cells died in G1/S transition or S phase, characterized by an elongated cell shape and a small bud (Figure 1A and B for definition and some examples). No death is assigned to G2 as it is not obvious in budding yeast.

**Figure 1.**
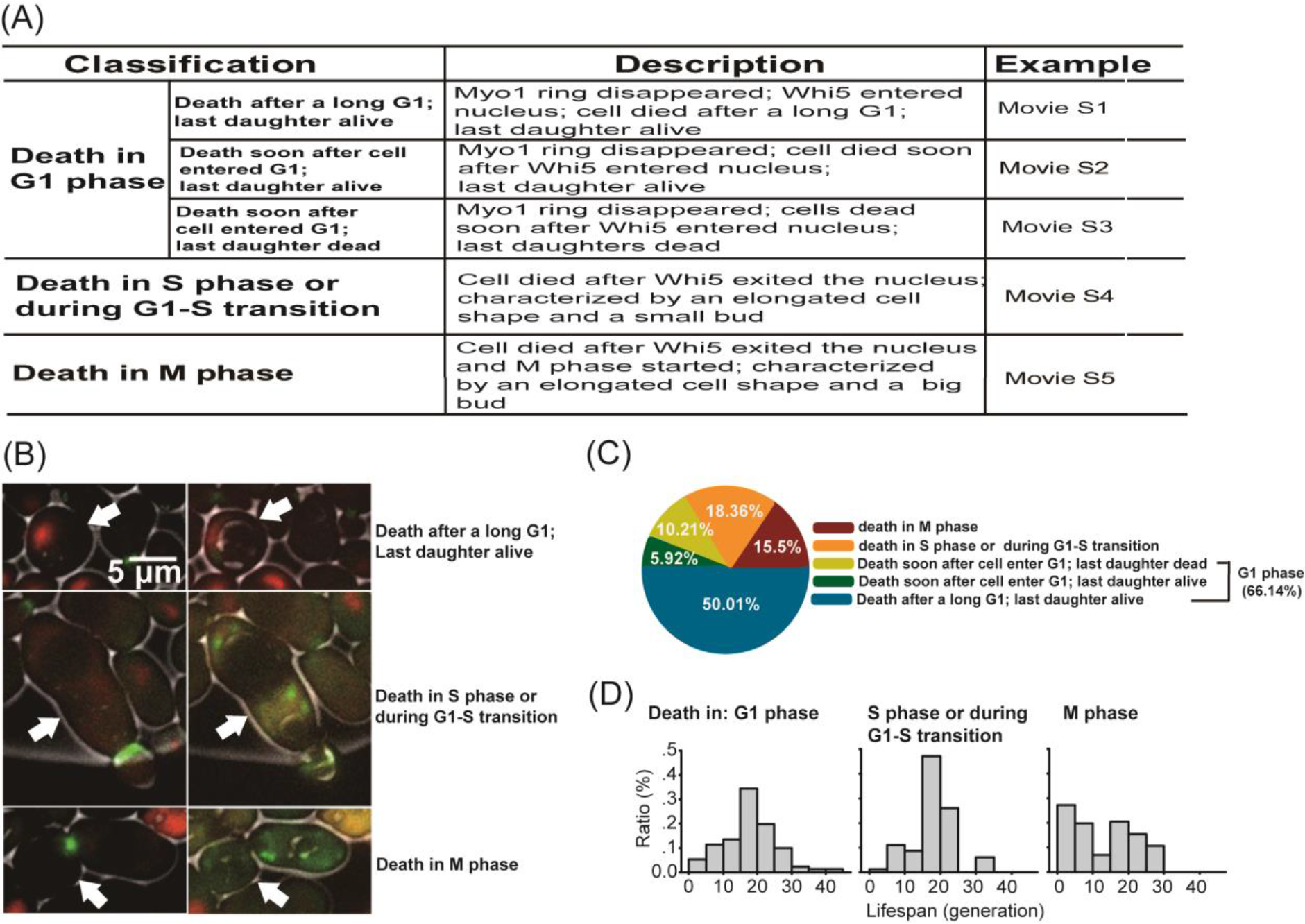
Cell death in different cell cycle phases and the corresponding lifespan distributions. (A) Classification and description of cells died in different cell cycle phases. (B) Bright field and fluorescent images (green: Myo1-GFP; red: Whi5-tdTomato) of cells died in G1 (top), S or G1-S transition (middle), and M phase (bottom). Left and right columns are images before and after death. White arrows indicate the mother cells. (C) Distribution of cells died in different cell cycle phases. The total cell number of cells is 204. (D) Distributions of replicative lifespan for cells died in different cell cycle phase.

We found that the majority of the cells died in G1 phase, consistent with the previous analysis based on terminal morphology using the traditional micro-dissection technique (Delaney, Chou et al. 2013). Among a total of 204 cells we analyzed, 66% died in G1 phase, 16% died in M phase, and the rest of 18% died in S phase or during G1/S transition. The 66% G1 death can be further divided into three different categories based the length of the stay in G1 and the fate of the daughter cell: 50% died after a long G1 with the last daughter viable, 6% died soon after entering G1 with the last daughter viable, and 10% died soon after the cytokinesis is finished (Myo1 ring disappeared) and the last daughter cell died too (Fig. 1A-C and Movies S1-S5), possibly due to chromosome damage from the last cell division.

We next measured the lifespan distribution for cells died in different cell cycle phases and found distinct characteristics. For G1-death and S-death, the corresponding lifespan distributions have a bell shape, with a characteristic age (peak of the distribution) close to the median lifespan (~20 generations). In contrast, for cells with M-death, the lifespan distribution is flat with no characteristic age (Fig. 1D). This indicates that for the former, there is very little death in young cells and the rate of death starts to be significant around the median lifespan (where rate of death multiplied by the number of live cells reaches a peak). For the latter, cell death happened significantly in young cells and throughout different ages. We hypothesize that G1-death resembles death due to normal aging while M-death resembles accidental death possibly caused by errors during chromosome segregation.

We found that cells died in M phase have significantly shorter lifespan than those died in other phases (G1, G1-S/S), with the median lifespan of 15 compared to 20 for the later (p-value<0.005, Chi-Square test). This results from significant death in young cells and is consistent with the previous observation that cell died with budded morphology have shorter lifespan compared with those died with un-budded morphology (Delaney, Chou et al. 2013). Cells with M-death also have broader lifespan distribution, consistent with the bigger variance of lifespan distribution for cell died with budded morphology (Delaney, Chou et al. 2013).

### Cells with Gl-death have prolonged cell cycle in the last few cell divisions mainly due to the lengthening of G1 phase

Since the majority of cells died in G1 and we hypothesized that G1-death represents death due to normal aging, we further analyzed how the cell cycle length and G1 length change over the course of aging for cells died in G1. Cell cycle length is defined as the interval between two successive budding events and G1 length defined as the duration of Whi5’s stay in the nucleus (Fig. 2A-B, Movie S6). Through most of their lifespan, cells have a constant cell cycle length and G1 length. However, for the last few cell divisions (typically the last 2-3 cell divisions), both the cell cycle length and the G1 length increase significantly (Fig. 2C-D), and can reach hundreds of minutes.

We found that the prolonged cell cycle in the last few cell divisions is mainly due to the increase of G1 length. The long cell cycle in the last few divisions is linearly proportional to the G1 length, and can be fitted by the relation Cell_cycle_length = 1.12*G1_phase_length + 90 min (Fig. 2E). This implied that the increase of cell cycle length results mainly from the increase of G1 length.

**Figure 2.**
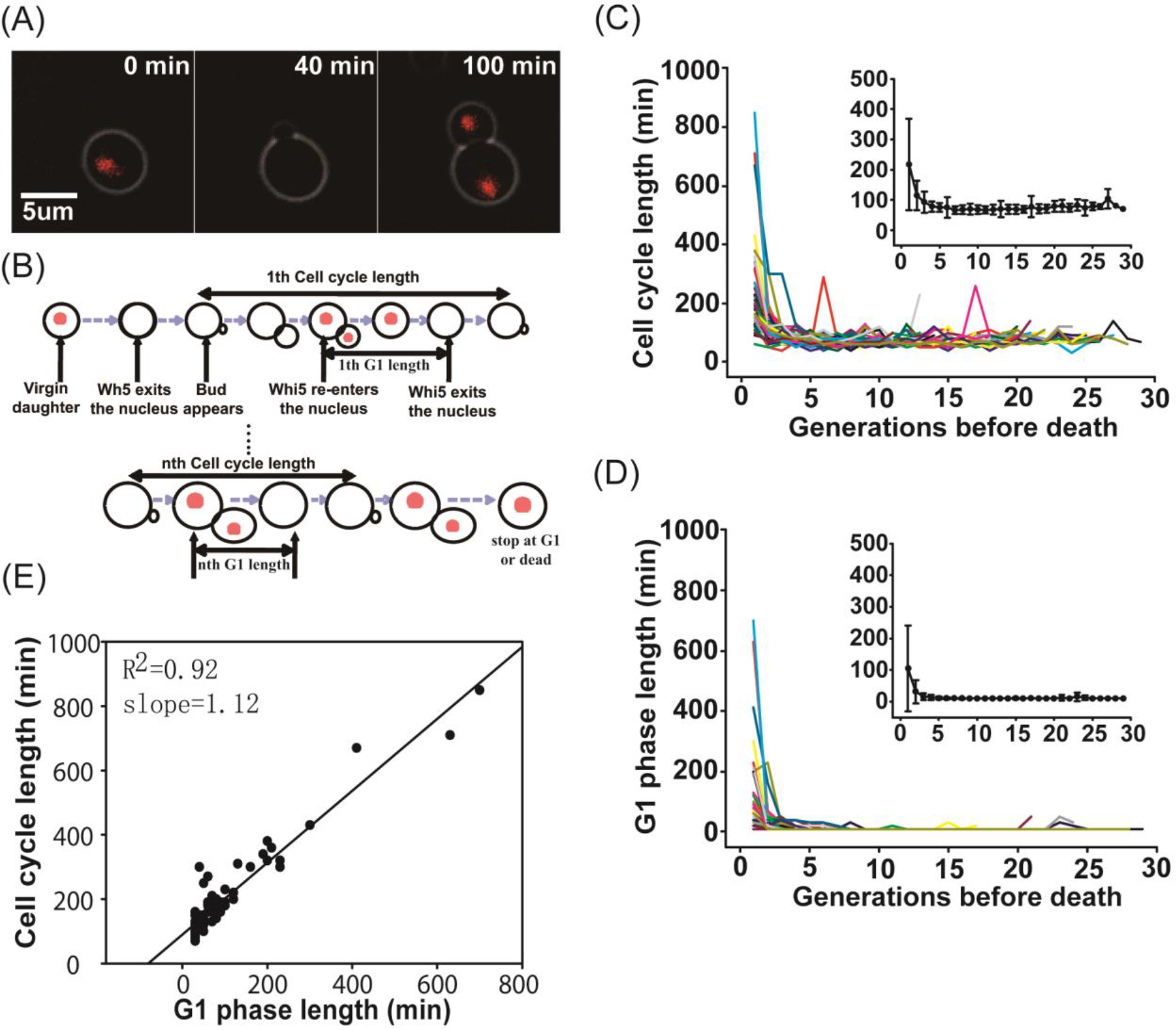
Cell cycle length and the duration of G1 phase as a function of age. (A) Composite phase contrast images showing Whi5 in and out of nucleus (red: Whi5-tdTomato). (B) A schematic on how cell cycle length and G1 length are measured. G1 length is defined as the duration of Whi5’s stay in the nucleus and cell cycle length is defined as the time interval between two successive budding events. (C) & (D) Cell cycle length and G1 length as a function of the generations before death in single yeast cells. Each colored line represents one single cell (total number of cells = 55). The average and standard deviation are shown in the corresponding insert. The measurements were done using 40 hours time-lapsed images (bright field and Who5-tdTomato) with 10 min interval. For cells with G1 shorter than 10 min, images of Whi5 in the nucleus may not be captured and the corresponding G1 length is taken to be 10 min. (E) G1 phase length vs. cell cycle length for cells died in G1. The linear fit is for data points with G1 length > 20 min.

There are a few exceptions to the general behavior described above. E.g., occasionally a cell can have a long cell cycle but short G1 in their middle life, but is able to recover and go back to normal cell division (Fig. 2C). This is diagnostic of problems in other cell cycle phases such as M phase, consistent with the notion that problems occurred in M phase early in life are caused by accident.

As aging is believed to be caused by cumulative damages that happen throughout life, the observation that aging impacts cell cycle only in the last few cell divisions suggest that the cell cycle control machinery respond to aging signal to alter G1 and cell cycle length only when the stress signals exceeds a certain threshold.

### Whi5 nuclear concentration increases with age and correlates with the length of G1 phase above a threshold concentration

The prolonged G1 phase before death in old cells prompted us to analyze the dynamics of Whi5 as a function of age. Whi5 is a key transcriptional repressor of G1/S transition and was previously reported to participate in size control (Avraham, Soifer et al. 2013). It has been proposed that in the newborn daughter cells the dilution of Whi5 during the pre-START G1 phase triggers START, serving as a mechanism to regulate G1 length and cell size (Schmoller, Turner et al. 2015). Given Whi5’s role in the START checkpoint and the regulation of G1 length in newborn daughter cells, we analyzed how Whi5 nuclear concentration changes as a function of age in single mother cells.

Using the strain with Whi5 and Myo1 markers, we monitored Whi5 nuclear concentration once every 10 minutes throughout the lifespan of mother cells (see Methods). As expected, Whi5 nuclear concentration shows one pulse per cell cycle, reaching a peak in early G1 (Fig. 3A,B). From young to middle age (the first ~10 generations), the peak concentration remains constant while the cell size steadily increases with time (supplementary Fig. S2). Starting from middle age (9~10 generations before cell death), Whi5 peak concentration gradually increases with age and reaches a high value, typically 3 to 5 fold higher than that in the young cells, 2-3 generations before death (Fig. 3C).

**Figure 3.**
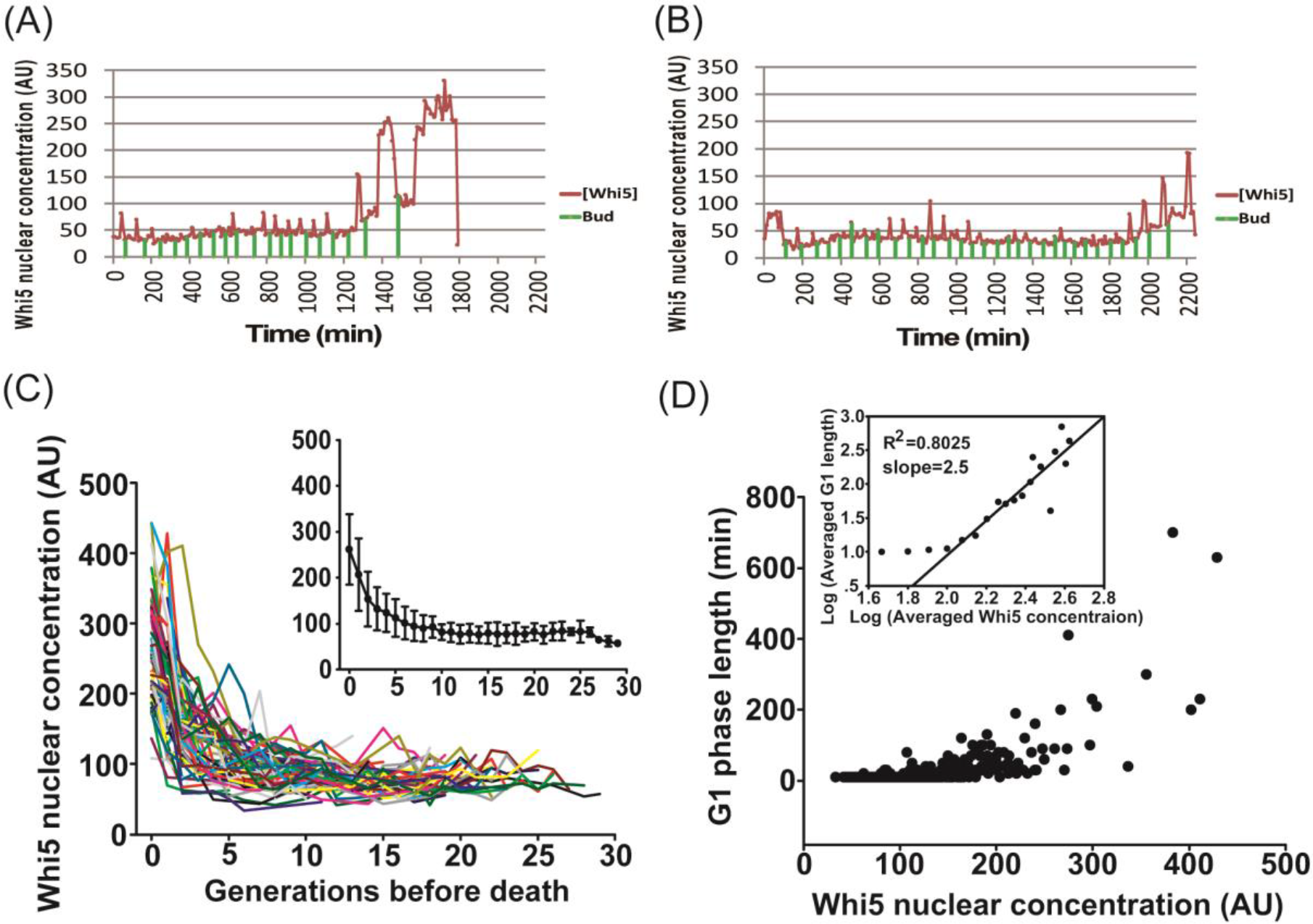
Whi5 nuclear concentration increases with age and is correlated with the length of G1 phase above a threshold. (A) & (B) Examples of Whi5 nuclear concentration as a function of time in single cells throughout their lifespan. 40 hours time-lapsed images were taken with 10 min interval. Green bars indicate the budding events. (C) Whi5 peak nuclear concentration as a function of generations before death in single cells. For cells with G1 length≥20 min (Whi5 in the nucleus in three or more consecutive frames), the highest Whi5 nuclear concentration is defined as the peak concentration. Each colored line represents one cell (total number of cells = 55). The mean and standard deviation is shown in the insert. (D) Correlation between Whi5 nuclear concentration and G1 length. Each point represents one pair of measured G1 length and Whi5 nuclear concentration (total number of cells = 55, total number of pairs =604). Insert: Log-Log plot of Whi5 nuclear concentration (binned with bin size=20) vs. the average G1 length in the bin. The linear fit is for data points with x≥2.07 (concentration≥120). The slope of the linear fit = 2.5.

We found that the drastic increase of Whi5 nuclear concentration and the significant lengthening of G1 phase near the end of the life is strong correlated. Focusing on the cells with G1 death, we measured the peak Whi5 concentration and the length of the corresponding G1 phase in each single cell throughout their lifespan. A scatter plot of G1 length vs. Whi5 peak concentration shows that they are correlated, but only above a threshold Whi5 concentration (~120) (Fig. 3D and the insert). Below the threshold, G1 length is independent of Whi5 concentration and assumes that of the normal G1 in young cells (about 10 min). Once above the threshold, G1 length increases with Whi5 concentration, with a faster than liner dependence (G1_length ~ [Whi5]^2.5 from the fit). This suggests that Whi5 nuclear concentration impacts the G1 length in aging cells only when it exceeds a threshold. Most of the cells reach this threshold only within their last 3-4 cell divisions (See Fig. 3A for examples and Fig. 3C insert, where the threshold concentration of ~120 mapped to 3-4 generations before death). This explains why significant lengthening of cell cycle (mainly G1) happens only in the last 3-4 divisions (Fig. 2C,D) while Whi5 concentration increases progressively starting from the middle age (Fig. 3C).

### Oxidative stress increases Whi5 concentration and slows down cell growth in a Whi5 dependent manner

To search for potential upstream signals that lead to the up-regulation of Whi5 during aging, we turned our attention to stress response. It has been observed that general stress response characterized by Msn2/4 activity increases progressively starting from the middle age (Xie, Zhang et al. 2012); the timing of which coincides with the increase of Whi5 nuclear concentration. It has also been shown that environmental stress can lead to the increased Whi5 activity through post-translational modification to suppress the expression of cyclins and slow down G1 progression (Gonzalez-Novo, Jimenez et al. 2015). We thus speculate that aging causes cellular stress due to the accumulation of molecular damages and/or loss of homeostasis and Whi5 expression increases in response to the stress. One natural candidate is oxidative stress, which has long been suggested as a major driver for aging (Harman 1956), and previous studies have observed increased ROS and oxidatively damaged proteins in old cells (Hanzen, Vielfort et al. 2016, Kaimal, Kandasamy et al. 2017).

To test whether Whi5 can respond to oxidative stress and slow down cell cycle progression, we treated yeast cells with different concentration of H_2_O_2_, and measured Whi5 total concentration and growth rate in the wild type and *WHI5* deletion mutant using flow cytometry (see Method). We observed a steady increase of Whi5 concentration over time after H_2_O_2_ treatment, in a dose-dependent manner (Fig. 4A). This is not due to the change of overall protein synthesis rate, or rate of dilution, as the other cell cycle marker Myo1 did not show response (Fig. 4B). H_2_O_2_ treatment significantly slows down cell growth in a dose-dependent manner, and Whi5 deletion significantly alleviates the growth inhibition by H_2_O_2_ (Fig. 4C). The rescue of the growth phenotype by *WHI5* deletion is partial, suggesting that there are redundant mechanisms to slow down cell cycle progression under oxidative stress.

**Figure 4.**
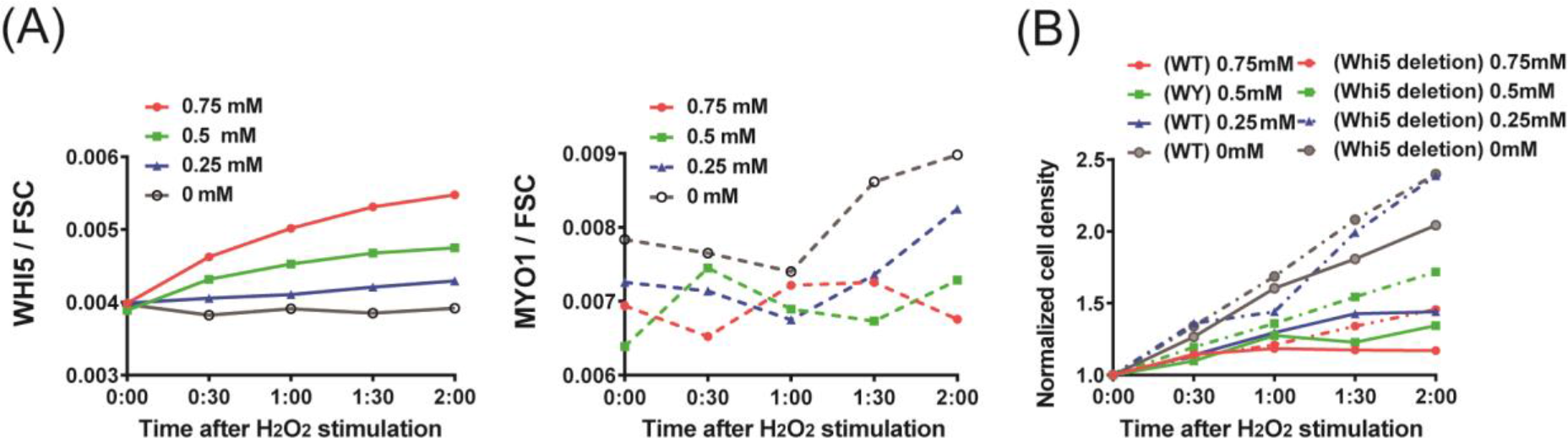
Oxidative stress increases Whi5 concentration and slows down cell growth in a Whi5 dependent manner. Cells were treated with different dose of H_2_O_2_ in their Log growth phase, and Whi5-tdTomato and Myo1-GFP were measured by FACS at different time points after the treatment. (A) Response of Whi5-tdTomato and Myo1-GFP to the treatments by H_2_O_2_ of different concentrations. Shown is the fluorescent intensity (total fluorescence normalized by the FSC for each cell) averaged over the cells. (B) Response of cell growth to H_2_O_2_ treatments in the wild type and the Whi5 deletion strains. Wild type refers to the Whi5-tdTomato and Myo1-GFP tagged strain.

### Whi5 deletion significantly extends yeast lifespan

The observed relation between stress, Whi5, and G1 length suggest a model that aged cells up-regulates Whi5 expression and arrests cell cycle in G1 in response to stress (Figure 6). This might be a conscious decision, i.e., arresting cells before the damage becomes too severe to allow cell cycle progression. This model makes the prediction that by removing Whi5, cells would delay such decision and therefore extends lifespan. Indeed, a previous study found that homozygous deletion of *WHI5* in diploid yeast cells (BY4743 background) extends the lifespan by about 30% (Yang, Dungrawala et al. 2011). We tested the lifespan of *WHI5* deletion in the haploid strain analyzed in this study and found that *WHI5* deletion increases the lifespan by about 40%, with a mean lifespan of 20.0 and 28.4 generations for the wild type and the *WHI5* deletion strains respectively (p =4.2*10^(−9), Fig. 5A). The averaged doubling times of the wild type and the *WHI5* deletion mutant are not significantly different, with 76.4 min and 75.9 min for the wide type and the mutant respectively. Interestingly, *WHI5* deletion also leads to a broader distribution of lifespan (Fig. 5B), suggesting increased stochasticity of cell death.

**Figure 5.**
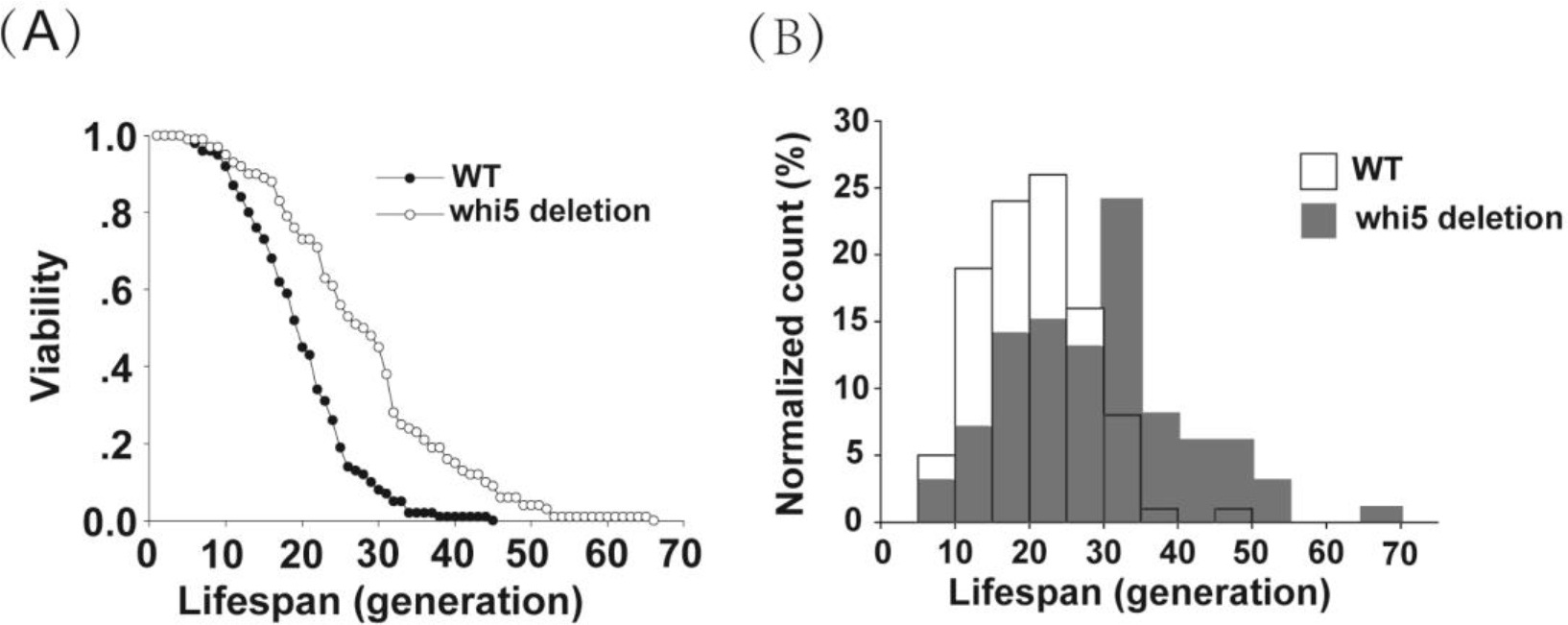
Whi5 deletion extends yeast replicative lifespan. (A) Survival curves of wild type and Whi5 deletion strain. (B) Whi5 deletion leads to a broader lifespan distribution compared to the wild type strain.

## Discussion

Yeast replicative aging ultimately leads to permanent cell cycle arrest. Understanding how and why cells respond to aging and stop cell cycle could lead to important insight into the mechanism. This has been elusive with the traditional micro-dissection technique, due to the lack of ability to continuously track mother cells and monitor molecular markers. Here by combining microfluidic devices with molecular tagging of important cell cycle regulators, we were able to analyze the cell cycle dynamics of single mother cells throughout their lifespan. We found that the majority of cells died in G1 phase, and their G1 phase lengthens considerably in the last few cell divisions before death. This lengthening of G1 correlates strongly with the nuclear concentration of Whi5, a major transcriptional repressor of G1/S transition. Interestingly this correlation is apparent only above a threshold concentration of Whi5; while Whi5 increases progressively starting from the middle age, G1 lengthening only occurs in the last few cell divisions after Whi5 reached a threshold concentration. We show that oxidative stress can lead to the increase of Whi5 expression and the slow down of cell division. Combining these data with the previous observations that general stress response (characterized by msn2/4 activity) starts to increase in the middle age, we propose a simple model that aging causes molecular damages/loss of homeostasis that activates stress response pathways to up-regulate Whi5, and to stop cell cycle at G1 once Whi5 reaches a high concentration (Fig. 6).

**Figure 6.**
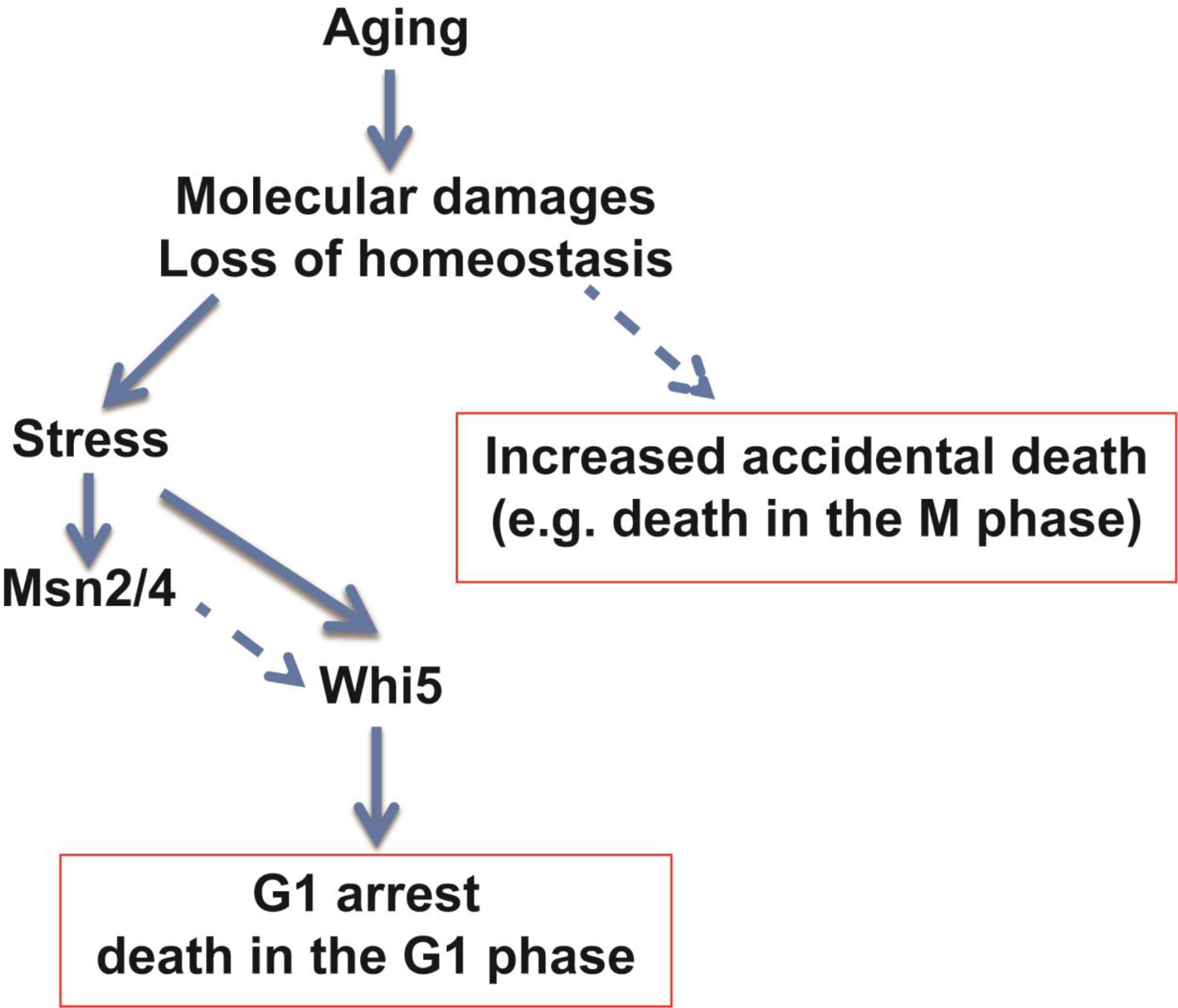
A simple model that connects aging with stress response and G1 death mediated by Whi5. Aging causes molecular damages and/or loss of homeostasis that activates stress response pathways to up-regulate Whi5, which eventually arrests cell cycle in G1 permanently. Solid lines indicate known connections and dotted lines hypothesized connections.

Stress response and its connection to cell cycle regulation have been studied extensively in cell cultures dominated by young cells. It is known that the general stress response regulators Msn2/4 respond to essentially all environmental stresses and up-regulate hundreds of genes to increase stress resistance and survival (Gasch, Spellman et al. 2000). Under high stress, cells typically arrest in G1 and re-enter cell cycle after the stress is removed or after they have adapted to the new environment. In a few case studies, the connection between stress and G1 arrest is known to be mediated by Whi5. For example, under osmotic stress, Whi5 nuclear retention can increase due to phosphorylation by Hog1 (Gonzalez-Novo, Jimenez et al. 2015), or due to the inhibition of Cln3-Cdc28 by Cip1, whose transcription is up-regulated by Msn2/4 (Chang, Tseng et al. 2017). Thus osmotic stress can communicate to Whi5 to slow down G1 progression through Msn2/4 dependent and independent pathways. Under oxidative stress, cells can arrest in G1 due to the oxidation of a specific cysteine residue in Swi6, which suppresses the expression of G1/S transition genes (Chiu, Tactacan et al. 2011). We showed that oxidative stress induced by H_2_O_2_ increases the total expression of Whi5 and slows down cell cycle (Fig. 4), suggesting that there is also a Whi5 dependent mechanism, and that the regulation of Whi5 is at transcriptional/translational level. We found that *WHI5* promoter contains the binding site for Msn2/4 (161 bps upstream) as well as several other stress response transcription factors including Hac1 (ER stress), Hsf1 (heat shock), and Yap1 (oxidative stress), suggesting that Whi5 is under transcriptional regulation by Msn/2 as well as regulators for specific stresses. Thus it seems that there are Msn2/4 dependent and independent mechanisms by which stresses can signal through Whi5 to cause G1 arrest (Fig. 6). The observation that over expression of Msn2/4 extends chronological lifespan (increased survival in the stationary phase) but shortens replicative lifespan (inhibiting budding) provided further genetic evidence supporting the links between Msn2/4, whi5, and G1 arrest (Fabrizio, Pletcher et al. 2004).

Whi5 nuclear concentration and Msn2/4 activity start to increase at middle age before any obvious cellular aging phenotypes are observed. Thus identifying the upstream stress signals that activate Msn2/4 and Whi5 may lead to important insight into the early molecular events that initiate the aging process. So far a number of candidates have been suggested (Lopez-Otin, Blasco et al. 2013), including oxidative damage, loss of proteostasis, decline of vacuole acidity (Hughes and Gottschling 2012), mitochondrial dysfunction, and dis-regulated nutrient sensing (such as aberrant TOR signaling). It will be informative to make reporters for Msn2/4 and Whi5 and one of the candidate stresses and analyze their dynamics of expression in single cells. Measuring the timing of different stresses relative to the increase of Msn2/4 and Whi5 may help identify the upstream signals. This could be an exciting direction to pursue in future studies.

Why does cell stop cell cycle at G1 in response to aging signals? The model we proposed here (Figure 6) suggests that the system is not evolved purposely to deal with aging, but instead evolved to handle stress and increase survival, and aging causes crosstalk to this pathway by inducing stress due to molecular damages and/or loss of homeostasis. Because of this crosstalk, mother cells permanently arrest their cell cycle in G1 “proactively”. Thus disabling Whi5 signaling can increase lifespan (Figure 5), but possibly at the cost of reduced fitness of the offspring and increased chance of chromosome abnormality.

We defined G1 as the period between Whi5’s entry to and exit from the nucleus. This definition of G1 has been adopted by several groups (Ball, Marchand et al. 2011, Liu, Wang et al. 2015) and can be more accurately measured than the traditional definition of G1, which refers to the period between the end of cytokinesis and the emergence of bud. The traditional G1 can be decomposed into two periods: *T1*, the period between the end of cytokinesis and Whi5 exit; and *T2*, the period between Whi5 exit and budding. *T1* is Cln3 and Whi5 dependent while *T2* is Cln1/2 dependent and cell size independent (Di Talia, Skotheim et al. 2007). Typically Whi5 enters nucleus a few minutes before the end of cytokinesis, as characterized by the disappearance of Myo1 ring (Figure 2B); thus the length of G1 defined by whi5 is close to *T1*. We showed that the lengthening of *T1* mostly accounts for the lengthening of cell cycle (Figure 2E), suggesting that it is the increase of Whi5 that is mainly responsible for the cell cycle lengthening, instead of the inability to induce the downstream Cyclins.

Our quantification of cell death in different cell cycle phases and the corresponding lifespan distributions is in concordance with the previous results based on terminal morphologies (Delaney, Chou et al. 2013), with small differences. E.g., we found that ~66% of cell died in G1, while Delaney et al. found ~78% of cells died with un-budded morphology, for the same BY4741 strain. We found that cells died in M phase have shorter lifespan and broader distribution, which is consistent with Delaney et. al.’s observation of cells died with budded morphology. However, the two different classifications are not exactly the same, e.g., a mother cell with a large round bud before the finish of cytokinesis may be classified as un-budded, while being classified as M phase death because the myo1 ring is still present.

Our analysis of the dynamics of Whi5 nuclear concentration revealed several interesting features suggesting important differences in cell size sensing and G1 control between newborn daughter cells and mother cells during aging. In newborn daughter cells, the initial concentration of Whi5 negatively correlates with the initial size, and the dilution of Whi5 through G1 growth triggers G1/S transition. Importantly, the rate of synthesis for Whi5 is independent of cell size while that for its upstream regulator Cln3 scales with cell size (size control mechanism, (Schmoller, Turner et al. 2015)). In mother cells, we observed a steady increase of cell size even in young cells (Fig. S2), while Whi5 nuclear concentration and G1 length is kept at constant. This suggests that the production of Whi5 is kept up with the cell size increase. Since the phosphorylation by the upstream Cdc28/Cln3 initially triggers Whi5 to exit, it is likely that the production of Cln3 also keeps increasing at the same rate as Whi5. From middle age to 2-3 divisions before death, Whi5 increases faster than cell growth but G1 length is still a constant. We speculate that during this period the production of Cln3 is kept up with the production of whi5 to keep G1 constant. This might also be a mechanism to impose a threshold response of G1 to stress. In the last 2-3 divisions, Whi5 increases beyond the threshold, perhaps no longer balanced by the increase of Cln3, to slow down G1 and eventually arrest the cell cycle permanently.

Cell cycle regulation in mammalian systems employs a parallel START check point, in which the transcription factor Rb plays a functionally analogous role as Whi5 in yeast, although the two factors bears no sequence similarity (Cooper 2006, Rowland and Bernards 2006). Rb is a negative regulator of cell cycle with an established role in cell senescence and is the first tumor suppressor identified (Cooper and Whyte 1989, Narita, Nunez et al. 2003). Our observations suggest that Rb may sense aging through stress response mechanisms and play an important role in establishing senescence in mitotically active cells during aging. With the inactivation of Rb, aging cells may continue to divide with increased chance of chromosome abnormality leading to cancer. The potential interplay between aging, stress, and cancer through Rb is a very interesting topic for future investigation (Chau and Wang 2003, Spike and Macleod 2005, Macleod 2008, Vurusaner, Poli et al. 2012, Salama, Sadaie et al. 2014).

## Methods

### Yeast strain and plasmid construction

Strains used for the experiments were derived from BY4741 (*MAT*a *his3 Δ 1 leu2 Δ 0 met15 Δ 0 ura3 Δ 0*). Myo1-GFP strain was taken from the yeast GFP library (Huh, Falvo et al. 2003). Whi5 deletion mutant strain was taken from the yeast deletion library (Giaever and Nislow 2014). To generate the strain with both Myo1-GFP and Whi5-tdTomato markers, we used the Myo1-GFP strain and tagged the endogenous Whi5 with tdTomato via homologous recombination, using the plasmid pCT2003 (*pRS306 WHI5pr-WHI5-tdTomato* (Liu, Wang et al. 2015)).

### Micro-fluidic chip preparation

Micro-fluidic devices used in the experiments were fabricated as described previously (Zhang *et al.*, 2012; Zou et al. JoVE 2016).

### Time-lapse microscopy and image analysis

For time-lapse microscopy, cells were prepared as previously described (Zhang *et al.*, 2012, Zou et al. JoVE 2016). Images were taken by a Nikon Ti-E time-lapsed microscope with Perfect Focus, Andor iXon3 897 EMCCD, Lambda SC shutter, Plan Apo VC 100×/1.4 oil DIC N2 objectives, and NIS-Elements Advanced Research software. A TOKAI HIT (INU-TIZHB-F1 Series) live cell incubator was used to keep the desired temperature and humidity. The micro-fluidic device was mounted on the microscope by a customized holder. Three sets of image were taken: 1) 8 hours of fluorescent images (Bright field/GFP/TRITC) with the interval of 3 minutes after 30 hours of bright field images with 10 minutes interval, to monitor cell death in different cell cycle phases (Fig. 1 & Fig. S1); 2) 40 hours of fluorescent images (Bright field/TRITC) with 10 minutes interval to measure G1 and cell cycle lengths, and Whi5 nuclear concentration (Figs. 2 & 3, and Fig. S2); 3) 60 hours of bright field images with 10 minutes interval to measure the RLS of *whi5 Δ* and wild type strains on the same microfluidic chip (Fig. 5). RLS was analyzed by counting the number of cell divisions of each mother cell from the time immediately after cell loading to cell death using the *ImageJ* software. Whi5-tdTomato fluorescent intensity in the brightest 9×9 pixel area was measured by *Image J*, and defined as Whi5 nuclear concentration. For long G1 divisions (G1 length ≥20), the highest intensity was defined as the Whi5 peak concentration of the corresponding cell cycle.

### Flow cytometry experiments for cells under H_2_O_2_ treatment

Yeast cells were cultured in tubes in a 30 °C shaker overnight and were diluted to OD_600_ = 0.1 next morning, and then grown in 96-wells plates for 3 hours, with 100 μl cell culture per well. Cells were then treated with different concentrations of H_2_O_2_ (0, 0.25, 0.5, and 0.75 mM). A BD Biosciences LSR II flow cytometer and HTS were used to measure the GFP and tdTomato signals [by the areas of the fluorescent isothiocyanate (FITC) channel (510 nm) and DsRed channel (586 nm), respectively] once every 30 minutes after the treatment (0, 30, 60, 90, 120min). To calculate the cell growth, cell density was calculated as previously described (Zuleta, Aranda-Diaz et al. 2014), with sample injection flow rate 0.5 μL/s and events number 20000/well.

### Normalization of data used for Figure 1

Data used for Figure 1 were taken by 8 hours of fluorescent imaging (Bright field/GFP/TRITC) with the interval of 3 minutes after 30 hours of bright field imaging with 10 minutes interval. We used 3 min interval fluorescent imaging to accurately determine the cell cycle phase in which a cell died. The fluorescent imaging was limited in the 8 hours window to avoid photo toxicity. We tracked cell death in this time window (to determine the cell cycle phase in which the cell died) and trace back by using the bright field images to count the RLS. This is different from the traditional lifespan assay (where all cells are tracked till death) and creates a small bias of the lifespan distribution (Fig. S1A). To correct for this bias we performed the following normalization: 1) count the RLS of the 112 cells by using the standard method (tracking all the cells till death, Zhang *et al.*, 2012). The lifespan distribution of these cells was used as a reference; 2) count the RLS of cells died in the 8 hours time window (number of cells=204) and generate the life span distribution (called Fig1_data); 3) Both reference data and Fig1_data were then divided into nine sub-groups based on the RLS, and a correction factor is calculated as the ratio of the frequency in Fig1_data vs. that in the reference data for each subgroup (Fig. S1B). The resulting correction factors were then used to correct for all the subsequent distributions shown in Figure 1. Fig. S1C showed an example of corrected lifespan distribution for cells died in M phase.

### Statistical Analysis

Comparisons of the lifespan distributions for cells died in different cell cycle phases were performed using Chi-square test in SPSS 17.0. Other statistic analyses were performed using Sigma plot 10.0, Sigma Stat 3.5 or GraphPad Prism7 software.

## Acknowledgments

We would like to thank Jonathan Weissman’s Lab for providing strains, and Zhengwei Xie, Ke Zou, Xuejun Zhu and other Ouyang Lab and Li Lab members for comments on the manuscript. This work was supported by a NIH grant AG043080, and a grant from the National Science Foundation of China NSFC81673338.

## Supplementary Information

**Fig. S1.**
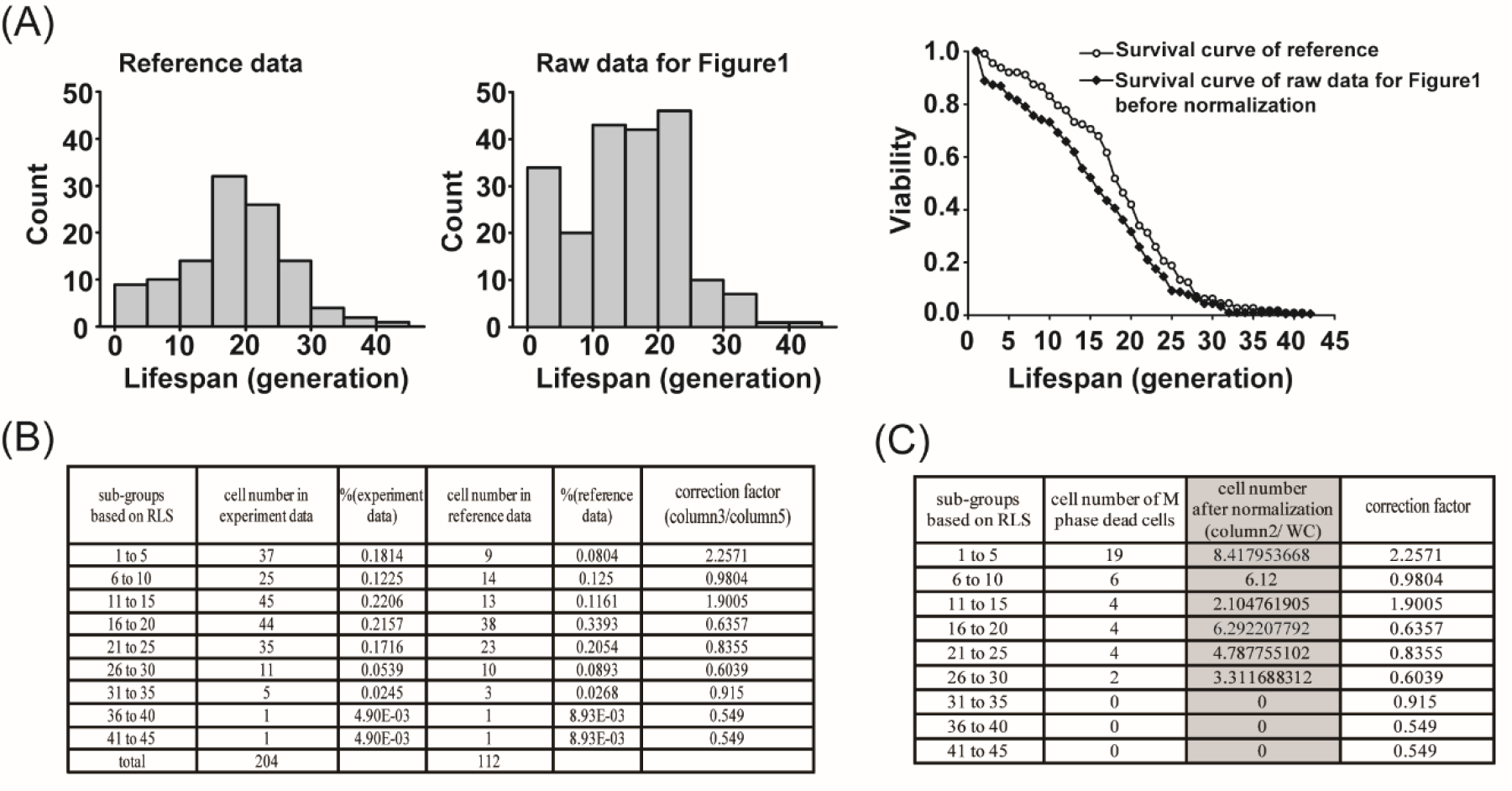
Data normalization for Figure 1. (A) The distribution and survival curve of RLS from the reference data (follow all the cells to death, cell number=112) and data used for Figure 1 (monitor cell death in a fixed time window, cell number=204). (B) Correction factors calculation for cells with different lifespan. (C) An example of lifespan distribution for cells death in M phase before and after normalization using the correction factors from (B).

**Fig. S2.**
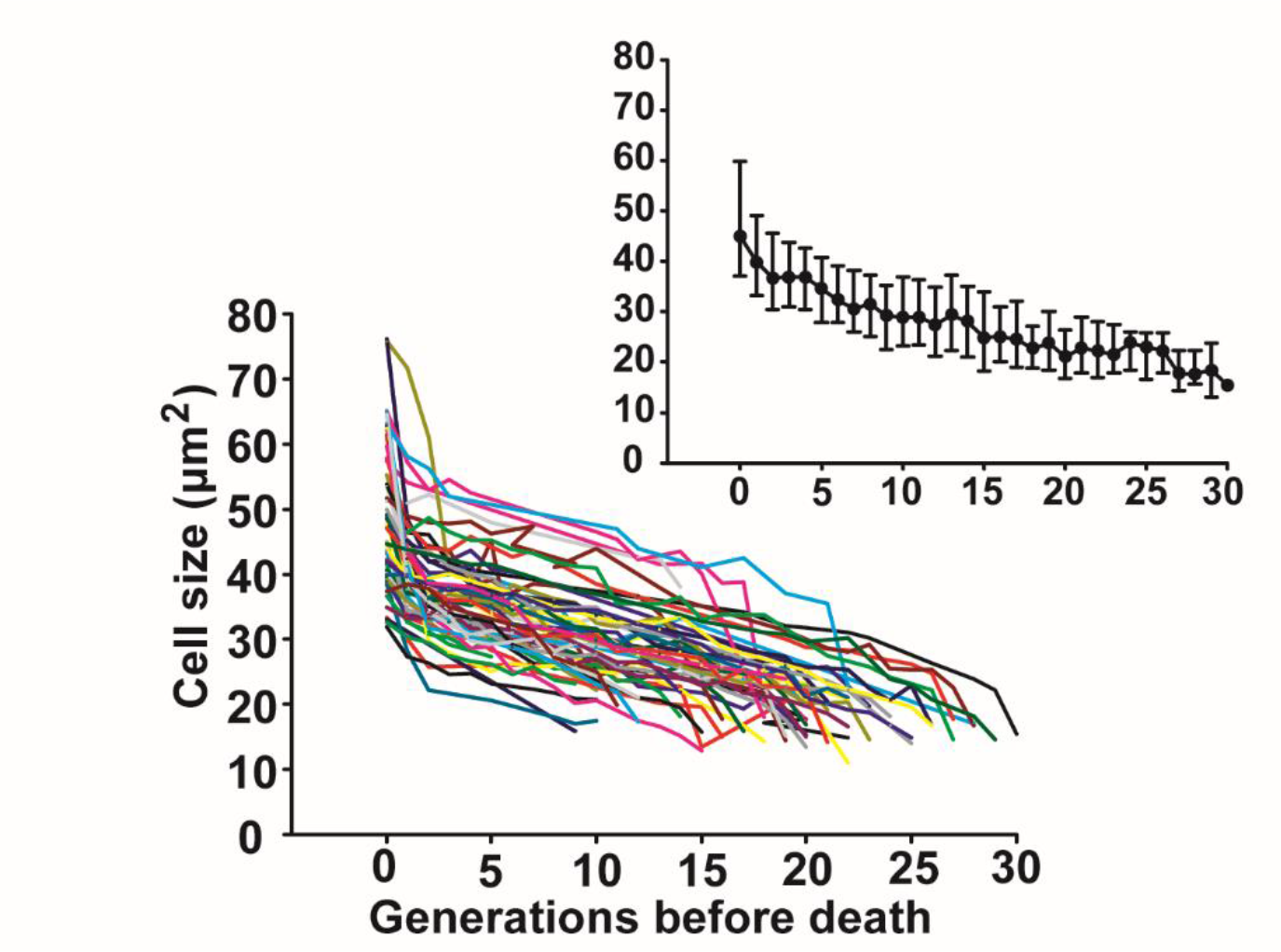
Cell size increases steadily with age. Cell size (measured by the surface area) is plotted against generations before death. Each colored line represents one single cell. The average and standard deviation are shown in the insert.

**Movie S1: A cell dead after a long G1 (RLS = 21 generations)**

**Movie S2: A cell dead soon after cell entered G1, last daughter alive (RLS=28 generations)**

**Movie S3: A cell Dead soon after it entered G1; last daughter dead (RLS=19 generations)**

**Movie S4: A cell Dead in S phase or during G1-S transition (RLS=20 generations)**

**Movie S5: A cell dead in M phase (RLS=24 generations)**

**Movie S6: Whi5 dynamics in aging**

**Additional data table S1 (separate file)**

Dataset S1: Raw RLS data for Figure S1 and Figure 1D

Dataset S2: Raw RLS data for Figure 5

